# Robust HIV-1 replication in the absence of integrase function

**DOI:** 10.1101/2020.03.18.997023

**Authors:** Ishak D. Irwan, Heather L. Karnowski, Hal P. Bogerd, Kevin Tsai, Bryan R. Cullen

## Abstract

Integration of the proviral DNA intermediate into the host cell genome represents an essential step in the retroviral life cycle. While the reason(s) for this requirement remains unclear, it is known that unintegrated proviral DNA is epigenetically silenced. Here, we demonstrate that HIV-1 mutants lacking functional integrase can mount a robust, spreading infection in cells expressing the Tax transcription factor encoded by human T-cell leukemia virus 1. In these cells, HIV-1 forms episomal DNA circles, analogous to Hepatitis B virus covalently closed circular DNAs (cccDNAs), that are transcriptionally active and fully capable of supporting viral replication. This rescue correlates with the loss of inhibitory epigenetic marks, and the acquisition of activating marks, on histones bound to unintegrated HIV-1 DNA. Thus retroviral DNA integration may have evolved, at least in part, as a mechanism to avoid the epigenetic silencing of extrachromosomal viral DNA by host innate antiviral factors.

**Significance:** While retroviral DNA is synthesized normally after infection by integrase-deficient viruses, the resultant episomal DNA is then epigenetically silenced. Here, we show that expression of the Tax transcription factor encoded by a second human retrovirus, HTLV-1, prevents the epigenetic silencing of unintegrated HIV-1 DNA and instead induces the addition of activating epigenetic marks, and the recruitment of NF-kB/Rel proteins, to the HIV-1 LTR promoter. Moreover, in the presence of Tax, the HIV-1 DNA circles that form in the absence of integrase function are not only efficiently transcribed but also support a spreading, pathogenic IN- HIV-1 infection. Thus, retroviruses have the potential to replicate without integration, as is indeed seen with HBV.

## Introduction

Integration of the proviral DNA intermediate into the chromosomal DNA of infected cells is a defining step of the retroviral life cycle (1, 2). Indeed, inhibition of integrase (IN) function is an effective means of blocking HIV-1 replication in not only T cells but also macrophages and IN inhibitors are routinely used as part of antiretroviral drug combinations (3, 4). However, the reason(s) why proviral integration is essential for productive HIV-1 replication remain unclear. Of note, even though hepadnaviruses such as hepatis B virus (HBV) also generate dsDNA copies of genome length viral RNAs by reverse transcription, chromosomal integration of this dsDNA does not form part of their replication cycle. So why is HBV able to replicate without chromosomal integration and, conversely, why is integration a critical step in all retroviral replication cycles?

A potential answer to these questions has emerged from the study of how not only HBV but also other DNA viruses, such as herpesviruses, are able to express their dsDNA genomes in infected cells. Specifically, HBx facilitates HBV gene expression by inducing the degradation of two cellular factors, SMC5 and SMC6, that otherwise block transcription of HBV episomes (5-7). SMC5 and SMC6 localize to PML nuclear bodies (PML-NBs), also called ND10, and depletion of PML-NB components, including PML and Sp100, rescues the transcription of HBV episomes even in the absence of HBx (8). PML-NBs also serve as components of the innate immune response to infection by several herpesviruses (9-11). In the case of herpes simplex virus 1 (HSV1), the incoming dsDNA genome is sensed by cellular factors, including IFI16, and then decorated with repressive chromatin modifications, including H3K9me^3^ and H3K27me^3^ (12-14). However, these inhibitory marks are then removed in a process that is dependent on the disruption of PML-NB function by the HSV1 immediate early protein ICP0 (12, 15). Thus cells have evolved an innate antiviral restriction pathway that recognizes extrachromosomal viral dsDNA as “non-self” and induces its epigenetic silencing (13). We propose that dsDNA viruses have evolved at least two mechanisms to counter this restriction pathway. In the case of HBV and most herpesviruses, this depends on viral proteins that induce the degradation of cellular factors that would otherwise epigenetically silence viral DNA, including SMC5/SMC6 in the case of HBV HBx, and the PML-NB components PML and Sp100 in the case of HSV-1 ICP0 (5, 11). Conversely, we hypothesize that retroviruses avoid cellular restriction factors that recognize and silence extrachromosomal DNA by integrating their proviral DNA into the host genome, where it eludes detection. It is well established that unintegrated retroviral DNA is rapidly loaded with histones that are then decorated with repressive chromatin marks that induce epigenetic silencing (16, 17). In the case of unintegrated murine leukemia virus (MLV) DNA, epigenetic silencing is mediated not by PML-NBs but rather by a DNA binding protein called NP220 acting in concert with the human silencing hub (HUSH) complex (18). However, the HUSH complex was reported to not play a role in silencing unintegrated HIV-1 DNA. Here, we report the surprising result that the epigenetic silencing of unintegrated HIV-1 DNA is prevented by ectopic expression of the Tax transcription factor encoded by a second human retrovirus, human T cell leukemia virus 1 (HTLV-1). In the presence of Tax, integrase deficient HIV-1 is able to effectively transcribe the circular DNA molecules that form in the nucleus in the absence of integrase function to mount a spreading, cytopathic infection.

## Results

If the inability of unintegrated HIV-1 DNA to be effectively transcribed is caused by host restriction factors, then the level of inhibition might vary between different cell types, as has in fact been previously reported (19). To confirm and extend this observation, we infected primary peripheral blood mononuclear cells (PBMCs) and various human cell lines with an NL4-3-based indicator virus in which the *nef* gene has been replaced with the Nano luciferase (*NLuc*) indicator gene (NL-NLuc). Cells were infected with WT HIV-1, with an IN mutant (D64V) that lacks integrase function, or with WT HIV-1 in the presence of 20 µM raltegravir (RAL), which blocks IN function (20, 21). Levels of NLuc expression were quantified and normalized to WT HIV-1, which was set at 100%. Similar levels of NLuc expression were observed whether IN activity was blocked by the D64V mutation or by RAL (Fig. 1A). These data revealed variable levels of inhibition of HIV-1 gene expression when proviral integration was blocked. Thus PBMCs, H9, CEM, CEM-SS and SupT1 cells all showed a >50-fold reduction in NLuc expression in the absence of IN function. Jurkat, HeLa, THP1 and A549 cells retained from 2% to 10% residual NLuc activity, while 293T cells allowed ∼12% residual NLuc expression in the absence of IN function. Remarkably, MT2 cells retained ∼70% of the NLuc expression in the absence of IN function while C8166 cells supported similar levels of NLuc expression whether IN was active or not (Fig.1A). Moreover, while infection of CEM-SS cells with the D64V IN mutant resulted, as expected, in minimal viral gene replication (Fig. 1C) and did not reduce cell viability (Fig. 1B), IN- HIV-1 was capable of almost WT levels of replication in C8166 cells (Fig. 1E), resulting in cytotoxic effects indistinguishable from WT (Fig. 1D).

**Fig. 1:**
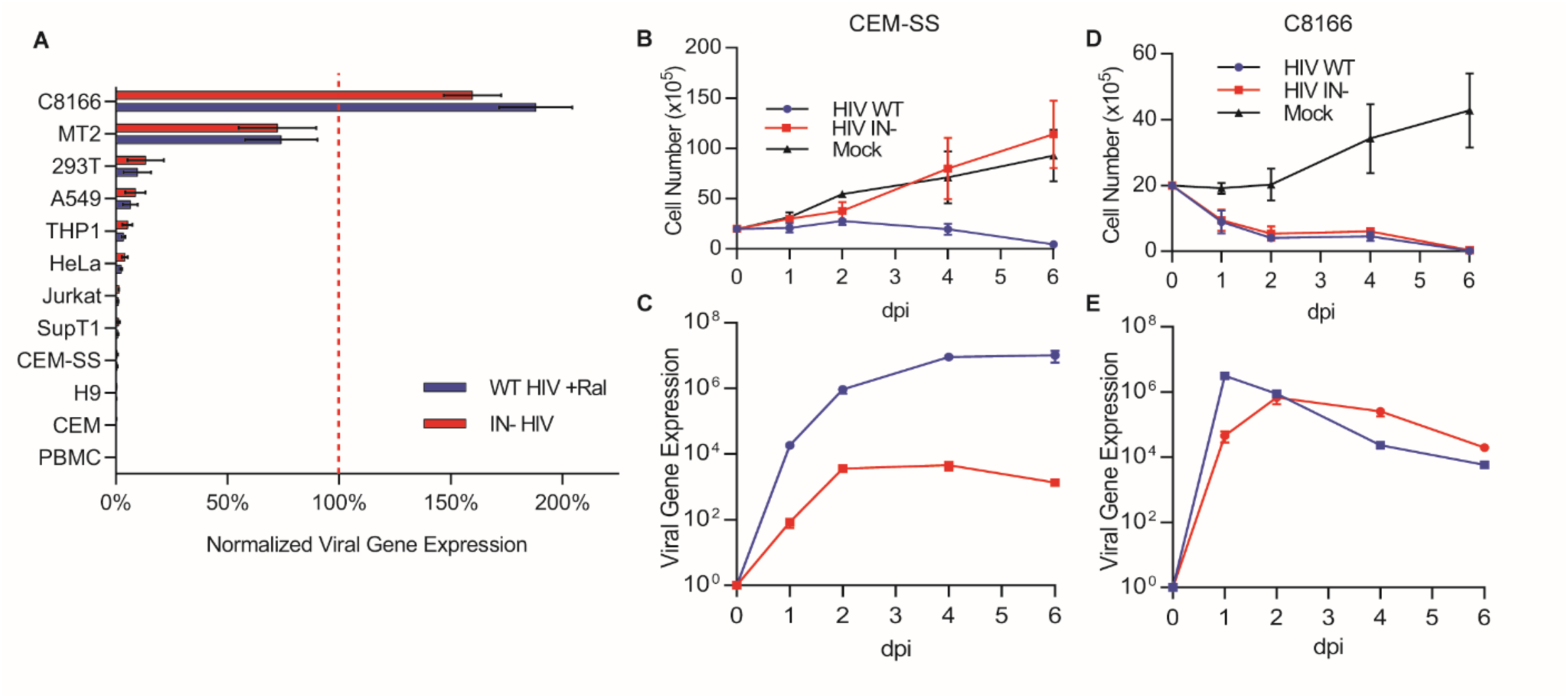
Differential gene expression and replication of IN- HIV-1. **A)** NLuc activity from the indicated cell lines or activated PBMCs infected with WT, WT+RAL, or with the D64V integrase mutant (IN-) NL-NLuc reporter virus at 48 hpi. The cells used express CD4 naturally or artificially. NLuc expression levels were normalized to WT, set at 100%. n=3, ± S.D. Viable cell counts of **B)** CEM-SS and **D)** C8166 cells infected with WT or IN- forms of the replication competent NL-NLuc indicator virus. Virally encoded NLuc expression was measured for **C)** CEM-SS and **E)** C8166 cells infected with WT or IN- NL-NLuc. All IN- infections in were conducted in the additional presence of 20 µM RAL.

Both C8166 and MT2 cells are HTLV-1 infected and express the HTLV-1 Tax protein, which can activate transcription from the HIV-1 LTR (22). To test whether Tax expression is sufficient to rescue the replication of IN- HIV-1, we transduced CEM-SS T cells with a Tet-inducible lentiviral vector expressing either Tax, or the HIV-1 Tat transactivator, the HIV-1 Vpr protein, which has been reported to partially rescue gene expression from IN- HIV-1 (23), or expressing the HIV-2 Vpx protein, which has been reported to block the inhibitory activity of the HUSH complex (24, 25). As shown in Fig. 2A, while expression of Tax fully rescued gene expression from IN- HIV-1 virus, it had little or no positive effect on the level of gene expression seen with WT HIV-1. In contrast, ectopic expression of the HIV-1 Tat protein had a small but non-specific positive effect on gene expression from both WT and IN- NL-NLuc, while expression of HIV-1 Vpr protein or HIV-2 Vpx both failed to affect either WT or IN- HIV-1 (Fig. 2A), although both proteins were expressed at readily detectable levels (Fig. S1B).

**Fig. 2:**
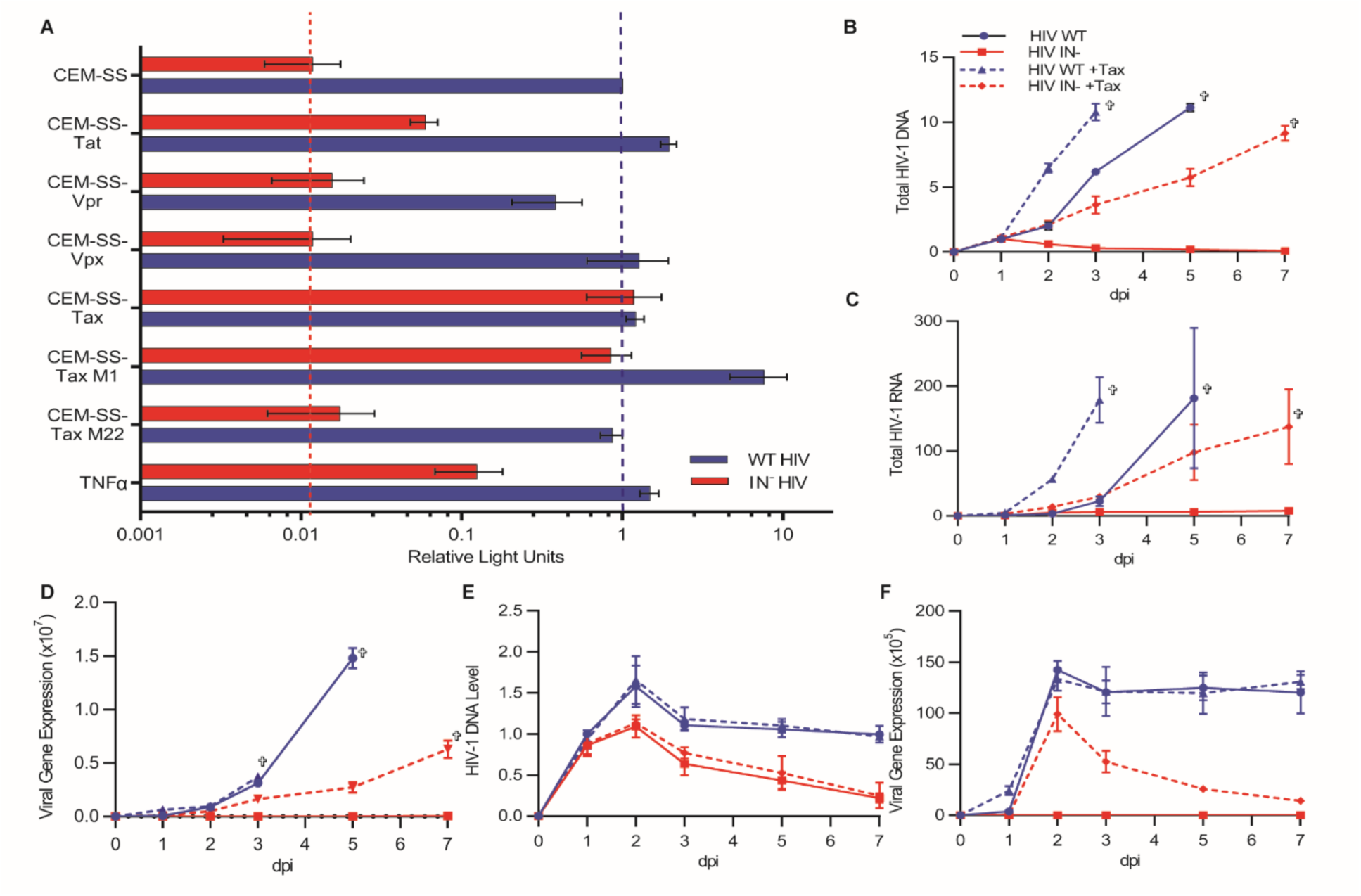
HTLV-1 Tax rescues integrase-defective HIV-1. **A)** CEM-SS cells were transduced with tet-inducible lentiviral vectors expressing each of the indicated viral proteins. Expression was induced with 0.5 µg/ml Dox and cells then infected with WT or IN- NL-NLuc. NLuc levels were measured at 48 hpi. TNF-α (1 ng/ml) was added at the time of infection. NLuc expression in each experiment was normalized to WT NL-NLuc-infected CEM-SS cells, which was set to 1. n=3, ± S.D. **B)** Clonal CEM-SS cell lines with tet-inducible Tax expression were infected with WT or IN- NL-NLuc reporter virus with and without Dox-induction. Total HIV-1 DNA levels in the cells were measured by qPCR at the indicated times. Similar to **B** but quantifying changes in **C)** total viral RNA, and **D)** Virally encoded NLuc expression over time. In **E** and **F**,Tet-inducible CEM-SS Tax cells were infected with non-spreading, VSV-G pseudotyped, IN+ or IN- NL-NLucΔEnv reporter virus in the presence and absence of Dox, and changes in **E)** total DNA, and **F)** NLuc expression quantified. All y-axes show fold changes relative to WT HIV-1 infected, uninduced (-Dox/-Tax) cells at day 1, which was set to 1; n=3. All IN- infections performed in the presence of 20 µM RAL. Crosses indicate last viable day of culture.

The HTLV-1 Tax protein can activate both the NF-kB pathway and the CREB/CRE pathway in expressing cells and Tax mutants lacking the ability to activate each of these pathways have been described (26). While the Tax M1 mutant, which lacks the ability to induce the CREB/CRE pathway, fully retained the ability to rescue IN- HIV-1 gene expression, the Tax M22 mutant, which is unable to activate NF-kB, lost the ability to rescue IN- HIV-1 gene expression, even though both mutants were expressed at similar levels (Figs. 2A and S1A). In further support of the critical role of NF-kB in this rescue, treatment of WT CEM-SS cells with TNFα, an inducer of NF-kB function (27), also selectively rescued gene expression from the IN- NL-NLuc vector, albeit not to the same degree as seen with Tax (Fig. 2A).

### Tax expression allows IN- HIV-1 to mount a spreading infection

Given that induction of Tax expression in the otherwise non-permissive CEM-SS T cell line can fully rescue IN- HIV-1 gene expression (Fig. 2A), we asked whether Tax could also rescue IN- HIV-1 replication. For this purpose, we analyzed the ability of WT or IN- versions of the otherwise replication competent NL-NLuc indicator virus to spread in CEM-SS cells transduced with the inducible Tax expression vector, in the presence and absence of Doxycycline (Dox). Parameters analyzed included total HIV-1 DNA levels (Fig. 2B), total HIV-1 RNA levels (Fig. 2C) and viral protein expression, as measured by analysis of the virally encoded NLuc protein (Fig. 1D) or of viral Gag expression (Fig. S1C). The IN- form of the NL-NLuc virus failed to express significant levels of viral DNA, RNA or protein in the absence of Tax while, in the presence of Tax, viral DNA, RNA and protein were not only readily detected but increased over the course of the experiment, resulting in the death of the CEM-SS culture by 7 dpi (Figs. 2B, 2C, 2D and S1C). To further confirm that Tax can indeed rescue the ability of IN- HIV-1 to mount a spreading infection, we repeated this experiment using a mutant of the NL-NLuc indicator virus lacking a functional *env* gene (ΔEnv) that cannot spread. WT and IN- forms of the NL-NLucΔEnv virus were pseudotyped with VSV-G and then used to infect CEM-SS cells with and without induced Tax expression. As shown in Fig. 2E, the ΔEnv IN- virus produced equal levels of HIV-1 DNA, regardless of Tax expression, that peaked at 2 days post-infection (dpi) and then gradually declined to background levels by 7 dpi, as predicted if the viral DNA was not chromosomally integrated. In contrast, while the ΔEnv version of the IN+ virus also gave rise to peak DNA levels at 2 dpi, there was only a modest decline in viral DNA by 7 dpi, regardless of Tax expression. While Tax had no effect on HIV-1 DNA levels in the context of a non-spreading infection, it did dramatically increase the level of viral gene expression from the IN- form of NL-NLucΔEnv, starting at 2 dpi. However, this positive effect was lost by 7 dpi, concomitant with the loss of unintegrated viral DNA (Figs. 2E and F).

To further confirm that Tax was not facilitating illegitimate HIV-1 DNA integration, we performed Alu-LTR qPCR (28), which quantifies the level of integrated viral DNA in HIV-1-infected cultures. As shown in Fig 3A, we observed high levels of integrated HIV-1 proviral DNA when the IN+ form of NL-NLuc was analyzed, while the IN- form of NL-NLuc, as predicted, gave rise to undetectable levels of integrated proviral DNA regardless of Tax expression.

**Fig. 3:**
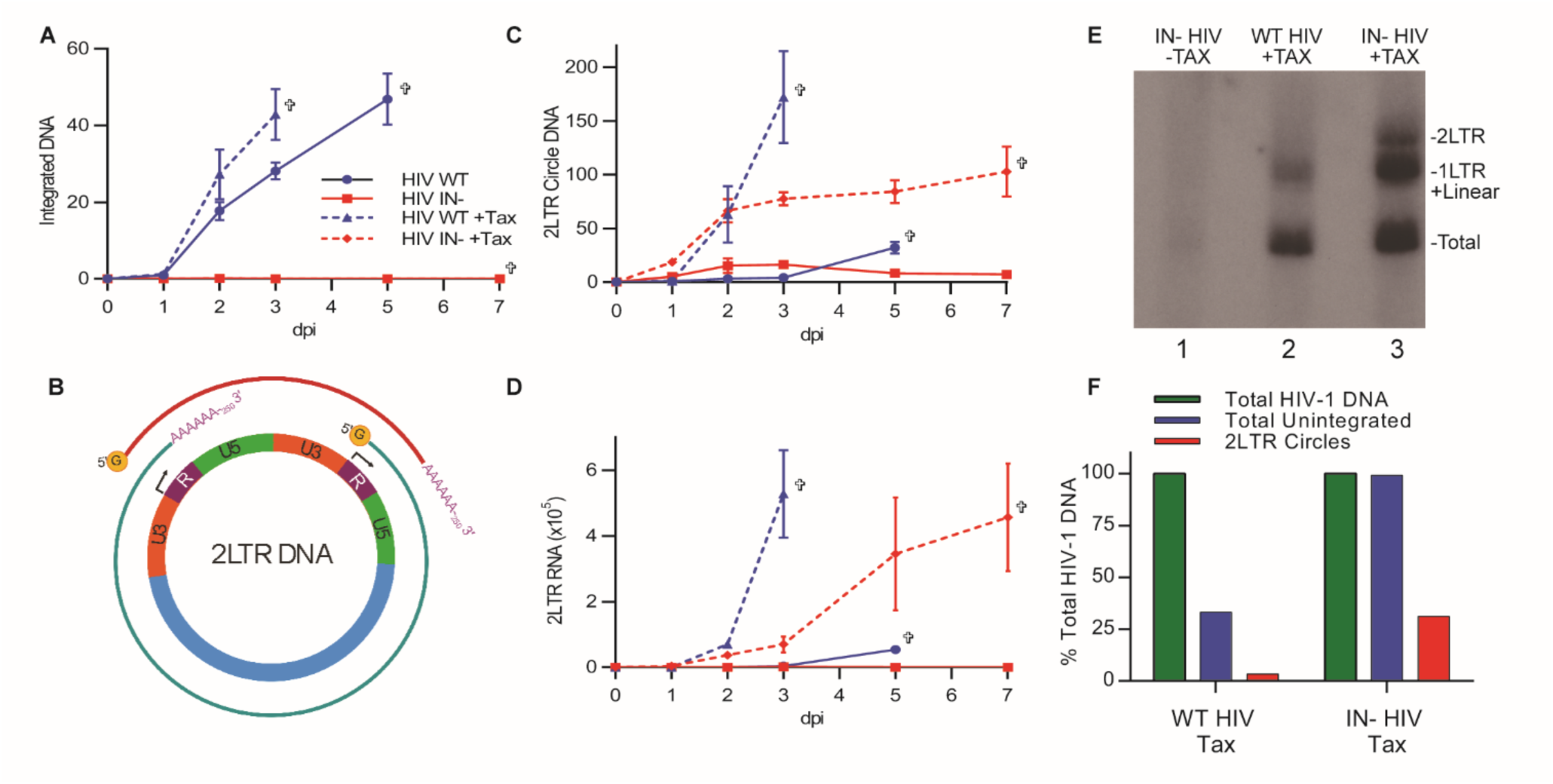
HTLV-1 Tax does not facilitate the illegitimate integration of IN- HIV-1. **A)** Quantification of integrated HIV-1 DNA in Tet-inducible CEM-SS Tax cells infected with WT or IN- HIV-1, in the presence or absence of Tax induction, as measured by Alu-LTR qPCR. **B)** Schematic showing the production of a short 2LTR transcript (red line) and the full-length transcript (green line) from circular 2LTR HIV-1 DNA. **C)** Total 2LTR circle DNA was measured at the indicated timepoints by qPCR, using primers that span the U5-U3 junction, in CEM-SS Tax cells infected with WT or IN- HIV-1 in the presence or absence of Tax. **D)** Quantification of the predicted 2LTR RNA transcript in CEM-SS cells ± Tax infected with WT or IN- HIV-1. All data in A, C and D are normalized to WT infected Tax-cells at day 1, set to 1; n=3. Crosses indicate last day viable cells were detected. **E)** DNA from CEM-SS cells ± Tax expression, infected with WT or IN- HIV-1, was digested with MscI and XhoI and probed on a Southern blot with HIV-1 probes that detect specific DNA species that correspond to 2LTR DNA (3.4 kb), 1LTR + linear unintegrated DNA (2.6-2.8 kb), and total HIV-1 DNA (1.9 kb). Note that lane 2 contains ¼ of the amount of cellular DNA used in lanes 1 and 3. See Fig. S2 for details. **F**) Quantification of the bands (2LTR+1LTR+linear and 2LTR) in **E** expressed as a percentage of total HIV-1 DNA, which was set at 100%.

### Integrase defective HIV-1 produces transcriptionally active DNA circles in the presence of Tax

Unintegrated HIV-1 DNA can exist in three different forms in infected cell nuclei. These are linear DNA, which is the substrate for integration; 1LTR DNA circles, formed by homologous recombination of linear DNA; and 2LTR DNA circles, formed by non-homologous end joining of the linear DNA (29). While all three forms of unintegrated viral DNA could be transcriptionally active, 2LTR circles are of interest because of their unique ability to generate a novel HIV-1 transcript, called 2LTR RNA, that initiates in what would normally be the 3’LTR and is then poly-adenylated in the 5’ LTR (Fig. 3B) (30). Analysis of the formation of 2LTR circular DNA in HIV-1 infected CEM-SS cells, in the presence and absence of Tax, revealed a low level in the IN- infected culture at early times after infection in the absence of Tax, but high and gradually increasing levels of 2LTR DNA in the presence of Tax (Fig. 3C). These data therefore mirror what was seen for total viral DNA in IN- virus infected CEM-SS cells in the presence and absence of Tax (Fig. 2B). Interestingly, Tax had an unexpected effect on the formation of 2LTR circles by IN+ WT virus. Specifically, while WT HIV-1 growing in CEM-SS cells in the absence of Dox induction gave rise, as expected, to low levels of 2LTR circles, this level was dramatically increased when Tax expression was induced (Fig. 3C). Similarly, when we analyzed the expression level of the predicted 2LTR RNA by qRT-PCR, we observed essentially undetectable levels in CEM-SS T cells infected by IN- HIV-1 in the absence of Tax but high and gradually increasing levels of 2LTR RNA when the same IN- virus was used to infect Tax expressing CEM-SS (Fig. 3D). Again, the WT IN+ version of the NL-NLuc indicator virus gave rise to only low levels of 2LTR RNA in the absence of Tax while the same virus, in the presence of Tax, produced high levels of 2LTR RNA (Fig. 3D). Thus, Tax causes WT HIV-1 to form a higher level of 2LTR circular DNA that is then efficiently transcribed to generate not only full length HIV-1 transcripts but also a short non-coding RNA consisting of two copies of the viral LTR (Figs. 3B and 3D). While the ability of Tax to induce transcription of the small number of unintegrated HIV-1 DNA proviral intermediates that arise during infection with WT HIV-1 is not unexpected, what was surprising was that Tax also dramatically increased the number of 2LTR proviral DNA circles (Fig. 3C).

To further confirm that Tax is indeed inducing a spreading infection due to the active transcription of unintegrated HIV-1 proviral DNA circles, we performed a Southern blot analysis to quantify total viral DNA, 2LTR DNA and linear/1LTR DNA (these latter two forms cannot be distinguished, see Fig. S2 for probe strategy). As shown in Fig. 3E, we again observed a dramatic increase in the level of HIV-1 DNA in Tax-expressing CEM-SS T cells infected with IN- HIV-1 (lane 3), when compared to the same virus in CEM-SS cells lacking Tax (lane 1). Moreover, when compared to IN+ HIV-1 (lane 2), quantification revealed that ∼100% of the viral DNA detected in the IN- infected culture was unintegrated, while ∼30 % of the viral DNA was unintegrated in the Tax-expressing culture infected with IN+ virus (Fig. 3F). Even though Tax at least modestly inhibits HIV-1 DNA integration in cells infected with WT HIV-1 (Fig. 3C), we only observed low levels of 2LTR circular DNA in these cells. In contrast, 2LTR circles contributed ∼30% of the viral DNA detected in Tax-expressing CEM-SS cells infected with IN- HIV-1 (Fig. 3F).

### Tax expression prevents the epigenetic silencing of unintegrated, chromatinized HIV-1 DNA

It has previously been reported that unintegrated HIV-1 DNA is rapidly loaded with core and linker histones and then epigenetically silenced due to the addition of inhibitory histone modifications (16). Therefore, we predicted that Tax, which induces the efficient transcription of unintegrated HIV-1 DNA, was likely acting to prevent this latter effect. We used ChIP-PCR to quantify the level of two inhibitory chromatin modifications (H3K9me3 and H3K27me3) and two activating chromatin modifications (H3K4me3 and H3Ac) on integrated and unintegrated HIV-1 DNA in the presence and absence of Tax. As previously reported (16), unintegrated HIV-1 DNA was significantly enriched for repressive H3K9 trimethylation, and depleted for activating H3K4 trimethylation, when compared to integrated proviruses (Fig. 4A). In contrast, in the presence of Tax, K9 trimethylation on H3 bound to unintegrated HIV-1 DNA was detected at levels that were not only significantly lower that seen with the same DNA in the absence of Tax but also lower than seen on integrated HIV-1 DNA (Fig. 4A). Similarly, the level of the activating H3Ac modification was significantly higher on unintegrated HIV-1 DNA in the presence of Tax than seen on either unintegrated or integrated HIV-1 DNA in the absence of Tax (Fig. 4A).We also observed a Tax-induced enhancement in the level of the activating H3K4me3 modification on unintegrated HIV-1 DNA, though this fell short of statistical significance (p=0.059). Finally, neither integration nor Tax expression affected the level of H3K27 trimethylation detected on viral DNA.

**Fig 4.**
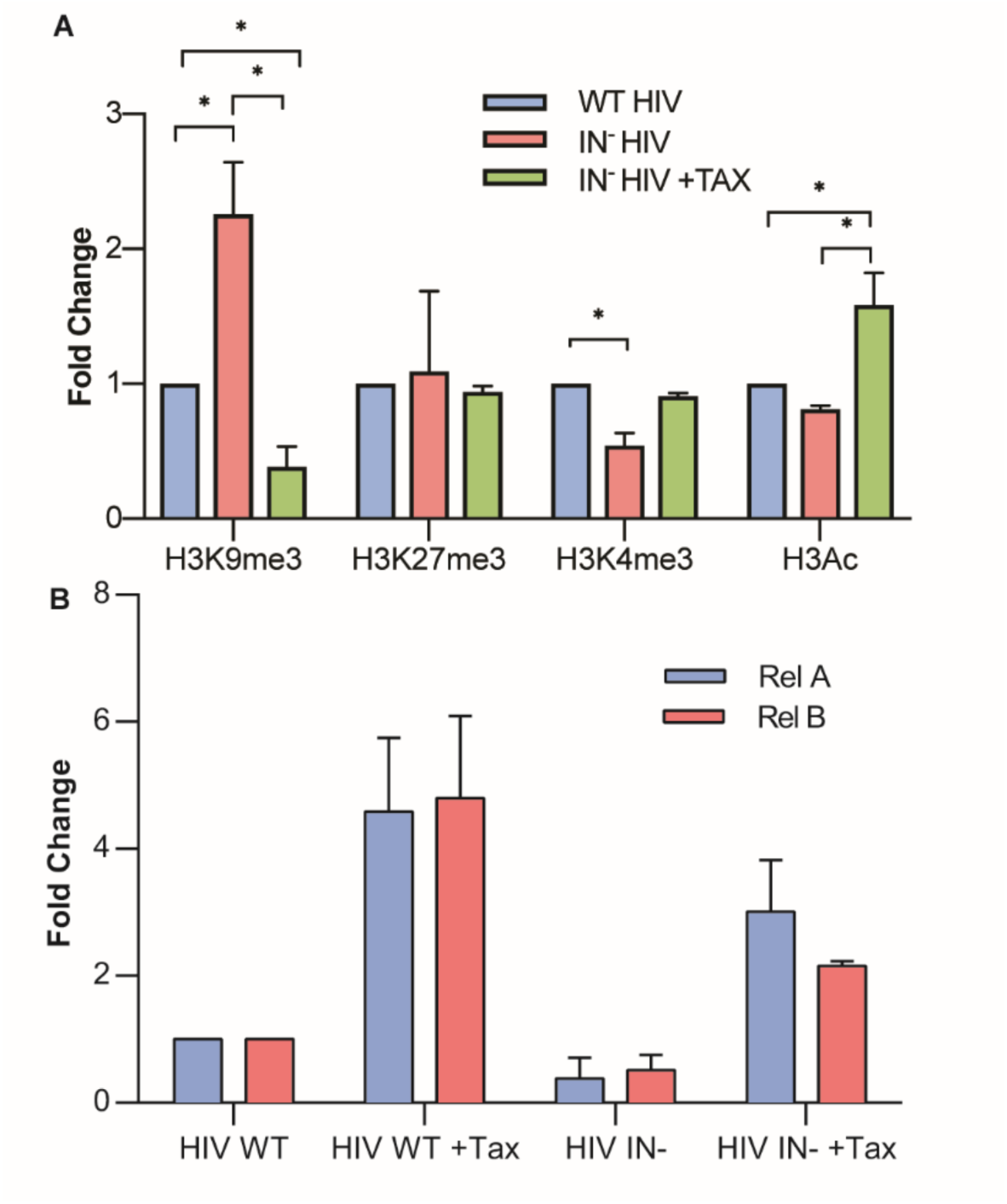
Tax induced changes in the epigenetic state and Rel/NF-kB binding at the LTR. **A)** CEM-SS cells in the presence or absence of Dox-induced Tax were infected with WT or IN- NL-NLuc and subjected to ChIP using antibodies against the indicated histone modifications. The level of bound HIV-1 DNA was quantified by qPCR, with WT IN+ HIV-1 set at 1.0. **B**) Similar to **A**, but Rel/NF-κB factors RelA and RelB were immunoprecipitated and bound DNA quantified by qPCR in CEM-SS cells ± Dox-induced Tax expression, infected with IN+ or IN- HIV-1. Cells were harvested at 2 dpi, and HIV-1 bound DNA quantified by primers that amplify part of the viral LTR. Fold changes were normalized to total histone H3 levels. n=3, ± S.D. * indicates p-values <0.05 (ANOVA with Tukey’s multiple comparison test). All changes in **B)** are statistically significant (p<0.01).

In Fig. 2A, we showed that the ability of Tax to rescue gene expression from unintegrated HIV-1 DNA not only required the ability to activate NF-kB but also that this effect could be partly mimicked by using TNFα to activate NF-kB. Previously, Tax has been reported to activate both RelA/p65 and RelB (31), both of which are potent transcriptional activators that are known to bind to the two canonical NF-kB DNA binding sites found in the HIV-1 LTR U3 region. We therefore performed ChIP-PCR to test whether Tax expression induced the recruitment of increased levels of RelA and/or RelB to the HIV-1 LTR. As shown on Fig. 4B, we indeed observed a large and statistically significant (p<0.01) increase in the recruitment of both RelA and RelB to both integrated and unintegrated HIV-1 DNA upon expression of Tax in infected CEM-SS cells.

As noted above, it has previously been reported that the epigenetic silencing of unintegrated MLV proviral DNA is mediated by the host cell DNA binding protein NP220 acting in concert with the HUSH complex (18). Surprisingly, however, this report also documented that the HUSH complex was not involved in silencing unintegrated HIV-1 DNA, a result that is consistent with the inability of the HIV-2 Vpx protein, a known inhibitor of HUSH complex function, to rescue gene expression from unintegrated HIV-1 DNA (Fig. 2A). We used CRISPR/Cas to mutationally inactivate the NP220 gene in human 293T cells by deletion of the entire NP220 DNA binding domain and simultaneous introduction of stop codons (Fig. 5A, Fig, S3A and B). Surprisingly, we observed that loss of NP220 function had no effect on the level of silencing of unintegrated HIV-1 DNA (Fig. 5B), thus further confirming that the silencing of unintegrated MLV DNA and unintegrated HIV-1 DNA are, in fact, mechanistically distinct. The obvious other potential cellular factor mediating unintegrated HIV-1 DNA silencing are PML-NBs, given their known role in the epigenetic silencing of other nuclear DNA viruses (10). However, knock down of several key PML-NB components, including PML, ATRX and Daxx, either individually or simultaneously, using RNA interference (RNAi) did not significantly enhance gene expression from unintegrated HIV-1 DNA (Fig. 5C and D). It could be argued that RNAi does not fully deplete targeted proteins and that the low, residual level of PML-NB function might still be sufficient to silence unintegrated HIV-1 DNA. To address this issue, we used CRISPR/Cas to inactivate the *pml* gene in 293T cells by introduction of frame shift mutations (Fig. S3C). However, loss of PML expression due to gene editing (Fig. 5E) also failed to rescue gene expression from unintegrated HIV-1 proviruses (Fig. 5F). Therefore, while the cellular mechanism that silences unintegrated HIV-1 DNA, and that is in turn inhibited by Tax expression, remains to be determined it does not appear to involve either the HUSH complex or PML-NBs.

**Figure 5:**
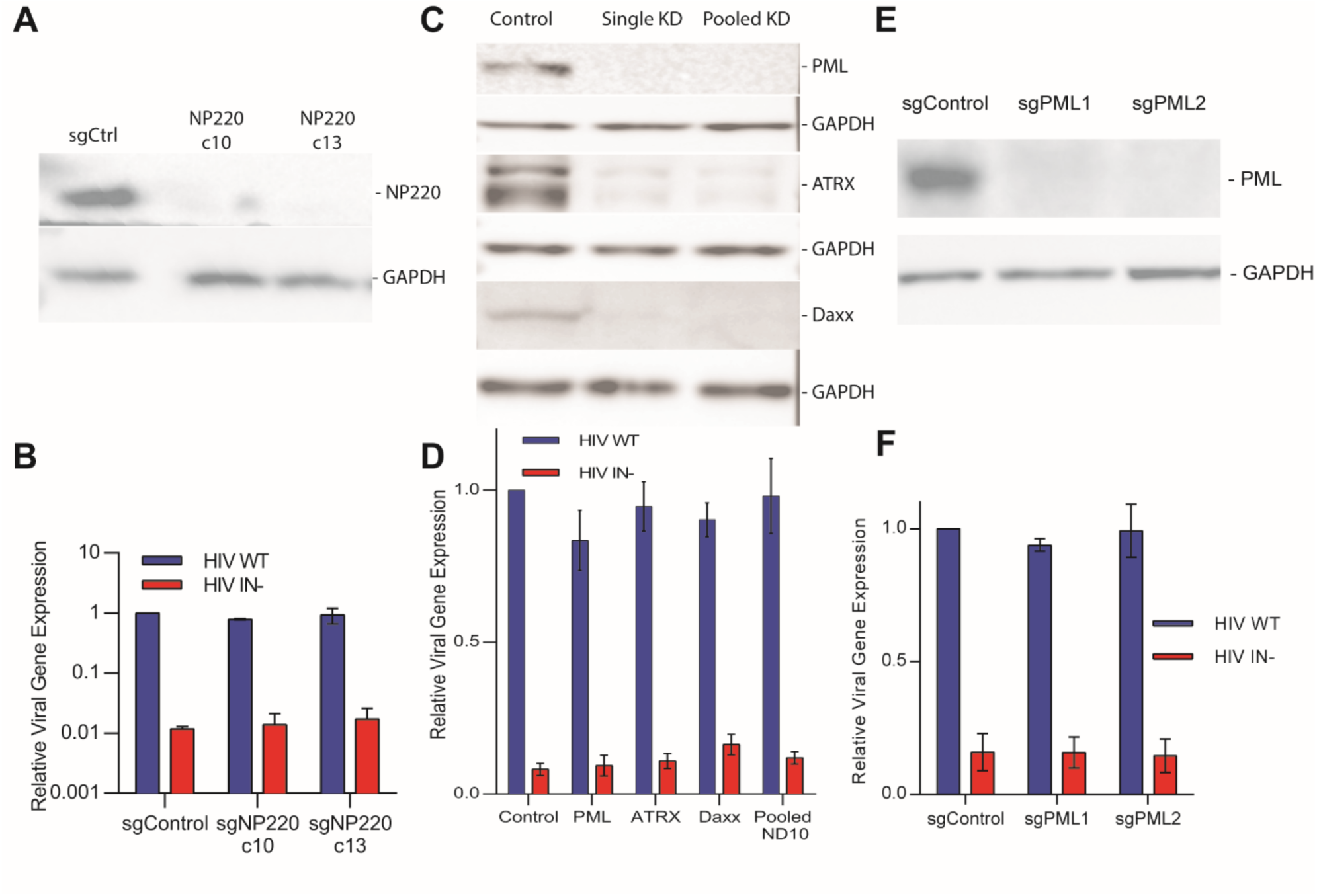
Neither NP220 nor PML-NBs appear to be involved in silencing unintegrated HIV-1 episomal DNA. **A)** We generated two 293T-derived clonal cell lines, c10 and c13, in which expression of NP220 was blocked by CRISPR/Cas-mediated gene editing. Loss of detectable NP220 expression was confirmed by Western blot. See Fig. S3 for details. **B)** The NP220 knockout 293T clones c10 and c13 were infected with VSV-G pseudotyped WT or IN- NL-NLucΔEnv reporter virus. NLuc expression was determined at 48 h post-infection and is given normalized to the IN+ indicator virus in control cells, which was set at 1.**C)** We used RNAi to knock down expression of the PML-NB components PML, ATRX or Daxx either individually (Single KD) or simultaneously (Pooled KD) in CD4-expressing 293T cells, as visualized by Western blot. **D)** The PML-NB component knockdown cells from panel C were infected with WT or IN- NL-NLuc virus, and NLuc expression determined at 48 h post-infection. Data are normalized to control cells treated with an irrelevant siRNA and infected with WT NL-NLuc, which was set at 1.0. **E)** Two CD4-expressing 293T clones, sgPML1 and sgPML2, were generated by knock out of the key PML-NB component PML using CRISPR/Cas. Loss of PML expression was confirmed by Western blot. See FigS3 for details. **F)** The PML knock out 293T clones sgPML1 and sgPML2 were infected with WT or IN- NL-NLuc virus, as described in panel B. NLuc expression was analyzed at 48 h post-infection and normalized to WT NL-NLuc in control cells, which was set at 1.0. For panels B, D and F, n=3 with S.D. indicated.

## Discussion

We report the surprising finding that expression in T cells of the HTLV-1 transcription factor Tax allows integrase deficient HIV-1 to mount a spreading, cytopathic infection characterized by increasing levels of HIV-1 DNA, RNA and gene expression (Fig. 2). These increases in viral DNA and protein expression were lost when a replication incompetent, env-deficient HIV-1 was analyzed, and hence must result from virus spread (Fig. 2E and F). Interestingly, while all the HIV-1 DNA produced during the spreading infection mounted by IN- HIV-1 in the presence of Tax is, as expected, unintegrated (Figs. 3A and 3F), much of this DNA is in the form of HIV-1 DNA circles, with 2LTR circles contributing ∼ 30% of the total (Figs. 3E and 3F). These 2LTR circles in turn produce readily detectable levels of a novel HIV-1 non-coding RNA consisting solely of the two viral LTRs (Figs. 3B and 3D). Unexpectedly, this novel 2LTR RNA was also detected in Tax expressing, but only minimally in non-Tax expressing, CEM-SS cells infected with IN+ HIV-1 (Fig. 3D), which correlated with a substantial increase in the production of 2LTR circular DNA in the Tax expressing cells (Fig. 3C). The reason for this phenomenon is not known but we speculate that Tax expression is activating cellular DNA binding proteins, including NF-kB (Fig. 4B), that bind to unintegrated HIV-1 DNA and induce the transcription of that DNA. This in turn may then sterically hinder proviral integration, resulting in an increase in transcriptionally active, unintegrated HIV-1 DNA even in IN+ HIV-1 infected, Tax expressing cells.

Previously, it has been reported that unintegrated retroviral DNA is rapidly chromatinized after nuclear entry and then decorated with inhibitory epigenetic marks that are removed, by an unknown process, after integration occurs (16, 17). Here, we demonstrate that Tax expression also reduces the deposition of inhibitory chromatin marks, and increases the deposition of activating marks, on unintegrated proviral DNA (Fig. 4A). This is turn correlates with an increase in the recruitment of the NF-kB/Rel family members RelA/p65 and RelB, both of which are activated by Tax, to the HIV-1 LTR on unintegrated HIV-1 DNA (Fig. 4B). Because activation of NF-kB is essential for the ability of Tax to rescue the viability of IN- HIV-1 (Fig. 2A), and because treatment with the NF-kB inducer TNFα also selectively rescues gene expression from infecting IN- HIV-1 (Fig. 2A), we speculate that NF-kB recruitment to the unintegrated viral DNA induces the observed epigenetic changes. However, Tax has also been proposed to play a role in chromatin remodeling in HTLV-1 infected cells (31, 32), so the mechanism of action of Tax may be complex.

Our data, and previous work from the Goff laboratory, are consistent with the hypothesis that proviral DNA integration allows escape from the epigenetic silencing that occurs if integration is blocked (17). However, this hypothesis does not preclude the possibility that integration also facilitates other aspects of retroviral replication. For example, integrated proviruses have the potential to be passed on to daughter cells after cell division and some retroviruses, such as HTLV-I, clearly use vertical transmission as a mechanism to increase viral load. In contrast, HIV-1 generally kills infected CD4+ T cells soon after infection while macrophages, the other major target cell for HIV-1, are non-dividing. In addition, in rapidly dividing cells, such as activated T cells, unintegrated HIV-1 proviruses, which lack an origin of replication, are rapidly lost (Fig. 2E). However, IN inhibitors also block HIV-1 gene expression in non-dividing macrophages (4, 33), so this cannot be the only reason for proviral integration.

Over 10 million people worldwide are infected with HTLV-1 and HTLV-1 infection is widespread in certain populations across the world, including in Australian aborigines (34). HTLV-1, like HIV-1, infects T lymphocytes, but unlike HIV-1, which rapidly kills infected T cells, the HTLV-1 Tax protein can activate cellular survival and proliferative pathways that instead induce T cell proliferation eventually leading, in a minority of infected patients, to an aggressive cancer called adult T-cell leukemia (31, 35). HTLV-1 infected CD4+ T cells can be fairly common in infected individuals, and the data reported here demonstrate that Tax expression can rescue the replication of IN- HIV-1 regardless of whether integrase function is lost due to mutagenesis if the *IN* gene or drug treatment (Figs. 1 and 2). We therefore speculate that used integrase inhibitors may be less effective in the treatment of dually HTLV-1 and HIV-1 infected individuals.

## Acknowledgements

This research was funded in part by a Duke University Center for AIDS Research (CFAR, P30-AI064518) pilot award to K.T. The following reagents were obtained through the NIH AIDS Reagent Program, Division of AIDS, NIAID, NIH: HIV-1 p24 Gag Monoclonal (#6458) from Dr. Michael H. Malim, HIV-1 NL4-3 Infectious Molecular Clone (#114) from Dr. Malcolm Martin, anti-Vpx monoclonal antobody (#2710) from Dr. John Kappes, CEM-SS cells (#776) from Dr. Peter Nara, MT-2 cells (#237) from Dr. Douglas Richmond and C8166 cells (#404) from Dr. Robert Gallo.

## Materials and Methods

### Key Resources Table

| Reagent or Resource | Source | Identifier |
| --- | --- | --- |
| Antibodies |  |  |
| Anti-H3K4me3 | EMD Milipore | cat#07-473, RRID: AB_1977252 |
| Anti-H3K27me3 | EMD Milipore | cat#07-449, RRID: AB_310624 |
| Anti-H3Ac | EMD Milipore | cat#06-599, RRID: AB_2115283 |
| Anti-H9me3 | Active Motif | cat#39161, RRID: AB_2532132 |
| Anti-Histone H3 | Abcam | cat#ab1791, RRID: AB_302613 |
| Anti-IgG | Cell Signaling | cat#2729S, RRID: AB_1031062 |
| Anti-p65 (RelA) | Santa Cruz | cat#sc-8008, RRID: AB_628017 |
| Anti-RelB | Santa Cruz | cat#sc-48366, RRID: AB_628212 |
| Anti-Tax | (26) | 6505 |
| Anti-Vpx | NIH AIDS Reagent | cat#2710 |
| Anti-Vpr | Proteintech | cat#51143-1-AP, RRID:<br>AB_10695191 |
| Anti-Tat | Hauber et al., 1987,<br>(36) | 705 |
| Anti-Gag | NIH AIDS Reagent | cat#6458, RRID: AB_2810883 |
| Anti-PML | Bethyl | cat#A301-167A-M, RRID:<br>AB_2779722 |
| Anti-ATRX | Sigma | cat#HPA001906, RRID:<br>AB_1078249 |
| Anti-Daxx | Sigma | cat#D7810, RRID: AB_259239 |
| Anti-NP220 | Bethyl | cat#A301-548A-M, RRID:<br>AB_1040051 |
| Anti-GAPDH | Proteintech | cat#60004, RRID: AB_2107436 |
| Anti-Mouse HRP | Sigma | cat#A9044, RRID: AB258431 |
| Anti-Rabbit HRP | Sigma | cat#A6154, RRID: AB_258284 |
| Anti-CD28/CD49d | BD Biosciences | cat#347690; RRID: AB_647457 |
| Bacterial and Viral Strains |  |  |
| HIV-1 isolate NL4-3 | NIH AIDS Reagent | cat#114 |
| NL-NLuc WT | Mefferd et al., 2018<br>(37) | N/A |
| DH5α | NEB | cat#C2988J |
| Chemicals, Peptides and<br>Recombinant Proteins |  |  |
| PEI | Polysciences | cat#23966-2 |
| Puromycin | Gemini Bio-Products | cat#400-128P |
| Doxycycline | Sigma | cat#D3447 |
| TNFα | Sigma | cat#H8916-10UG |
| WesternBright ECL kit | Advansta | cat#K-12045-D50 |
| Lipofectamine RNAiMax | Thermo | cat#13778150 |
| 37% Formaldehyde | Sigma | cat#252549-100ML |
| Critical Commercial Assays |  |  |
| GoTaq Green PCR Master<br>Mix | Promega | cat#M7123 |
| Nano luciferase assay kit | Promega | cat#N1110 |
| Power Sybr Green PCR<br>Mastermix | Applied Biosystems | cat#4367659 |
| Taqman Universal PCR<br>Master Mix | Applied Biosystems | cat#4324018 |
| HIV-1 p24 Antigen Capture<br>ELISA | ABL | cat#5421 |
| Dynabeads CD4 | Thermo | cat#11145D |
| Dynabeads Protein G | Thermo | cat#10004D |
| Antibiotic-Antimycotic | Gibco | cat# 15240062 |
| Prime-a-Gene Labeling<br>System | Promega | cat# U1100 |
| <sup>32</sup>P-dCTP | Perkin Elmer | cat#NEG013H100UC |
| Experimental Models: Cell<br>Lines |  |  |
| 293T | ATCC | cat#CRL-11268, RRID: CVCL_1926 |
| A549 | ATCC | cat#CCL-185, RRID: CVCL_0023 |
| HeLa | ATCC | cat#CCL-2, RRID: CVCL_0030 |
| THP-1 | ATCC | cat#TIB-202, RRID: CVCL_0006 |
| Jurkat | ATCC | cat#TIB-152, RRID: CVCL_0065 |
| H9 | ATCC | cat#HTB-176, RRID: CVCL_1240 |
| SupT1 | ATCC | cat#CRL1942, RRID: CVCL_1714 |
| CEM | ATCC | cat#CCL-119, RRID: CVCL_0207 |
| CEM-SS | NIH AIDS Reagent | cat#776, RRID: CVCL_J318 |
| MT-2 | NIH AIDS Reagent | cat#237, RRID: CVCL_2631 |
| C8166 | NIH AIDS Reagent | cat#404, RRID: CVCL_1099 |
| 293T-CD4 | Courtney et al., 2019<br>(38) | N/A |
| Oligonucleotides |  |  |
| All oligonucleotides are listed in Table S1 | This study | N/A |
| Recombinant DNA |  |  |
| pLentiCrispr v2 | Sanjana et al., 2014<br>(39); Addgene | cat#52961 |
| ΔCR8.74 | Addgene | cat#22036 |
| pMD2.G | Addgene | cat#12259 |
| pBabe-CD4 | Courtney et al., 2019<br>(38) | N/A |
| pTrex | Poling et al., 2017 (40) | N/A |
| Software and Algorithms |  |  |
| GraphPad Prism 8 | Graphpad Software | https://www.graphpad.com/scientific-software/prism/ |
| Image J | Rasband et al., 1997-2018 | https://imagej.nih.gov/ij/index.html |

### Cell lines used in this study

The human cell lines used in this study are 293T, an embryonic kidney epithelial cell line of female origin; CEM and CEM-SS are peripheral blood T lymphoblast cell lines of female origin; H9, SupT1, Jurkat, MT2, and C8166 are peripheral blood T lymphoblast cell lines of male origin; THP-1 is a monocytic cell line of male origin; HeLa is a cervical epithelial cell line of female origin; A549 is a lung epithelial cell of male origin. All cells were cultured at 37C with 5% CO2. 293T, HeLa, and A549 cells were cultured in Dulbecco’s Modified Eagle Medium (DMEM) supplemented with 10% fetal bovine serum (FBS) and Antibiotic-Antimycotic. CEM, CEM-SS, H9, SupT1, MT2, C8166, and THP-1 cells were cultured in Roswell Park Memorial Institute (RPMI) medium supplemented with 10% FBS and Antibiotic-Antimycotic. Primary CD4+ T cells were cultured in RPMI medium supplemented with 10% FBS, Antibiotic-Antimycotic and supplemented with IL2 72 h before infection.

CD4+ 293T, HeLa, and A549 cells were generated by transducing the cells with a retroviral vector expressing CD4 (pBabe-CD4), then selecting the cells for puromycin resistance. The cells were then single cell cloned, and the clonal lines screened for CD4 expression by infection with the NL-NLuc reporter virus, and assaying NLuc activity.

All Tet-inducible viral proteins were made in the modified lentiviral pTREX vector (40). HTLV-1 Tax, HIV-2 Vpx, HIV-1 Tat and HIV-1 Vpr were PCR amplified from existing expression plasmids, cloned into a modified pTREX vector and the sequence verified by Sanger sequencing. M1 and M22 Tax mutants were created by mutating the WT HTLV-1 Tax sequence (26) by overlap-extension PCR to introduce the relevant H3S (M1) or G137A/L138S (M22) mutations. Lentiviruses were packaged by transfecting 5 × 10^6^ 293T cells in a 15 cm dish with 15 µg of the lentiviral vector, as well as 10 µg and 5 µg of the packaging plasmids pCMVR8.74 and pMD2. G, respectively, using PEI. The media were changed 24 h post-transfection (hpt). Supernatants containing lentiviral particles were collected at 72 hpt, filtered through a 0.45 µM filter, and run through a 100,000 MWCO concentrator (Amicon). Following concentration, 5 × 10^6^ CEM-SS cells were incubated with 2 mL of the concentrated supernatant at 37 °C overnight. The media were then replaced with fresh RPMI medium and cells incubated for 48 h. At this point, the media were replaced with fresh RPMI medium supplemented with 1 µg/mL of puromycin to allow selection of transduced cells. Cells were then tested for Tet-inducible expression of viral proteins by Western blot.

### HIV-1 Production

For infections where NLuc was measured in spreading HIV-1 infections, we used an NLuc (pNL-NLuc) reporter virus in which the viral *nef* gene in NL4-3 had been substituted with the NLuc indicator gene (37). Non-spreading infections were similarly carried out using similar indicator viruses bearing a 943bp deletion in *env* and either NLuc (NL-NLucΔEnv), or GFP (NL-GFPΔEnv) in place of *nef*. These viruses either had WT integrase, or the D64V (IN-) mutation in the *integrase* gene that blocks function. Plasmids expressing the replication competent NL-NLuc provirus, or the non-spreading ΔEnv proviruses, were transfected into 293T using polyethylenimine (PEI). Non-spreading ΔEnv proviruses were co-transfected into 293T cells along with the pMD2.G plasmid encoding the VSV-G protein. After 24 h, the spent media were replaced with fresh media. At 72 hpt, supernatant media was filtered through a 0.45 mm filter. WT or IN- HIV-1 containing supernatant media were normalized by p24 levels, measured and normalized by ELISA, before being used to infect target cells.

### Purification of CD4+ T Cells from PBMCs

PBMCs were isolated from total blood by density gradient centrifugation (lymphocyte separation medium; Cellgro number 25-072-CV). CD4+ T cells were then isolated using the Dynabead CD4 positive isolation kit (Invitrogen; 1131D) following the manufacturer’s instructions. Cells were activated by incubation in phytohemagglutinin (PHA) and mouse monoclonal antibodies specific for human CD28 and CD49d (BD Biosciences) for three days, as previously described (41).

### Luciferase Assay

Cells were washed three times in PBS, lysed in Passive Lysis Buffer (Promega) and assayed for NLuc activity using the Nano-Glo Luciferase Assay on a Lumat LB9507 luminometer (Bertold Technologies).

### Analysis of HIV-1 Replication and Expression in Tet-inducible Tax cells

10^7^ Tet-inducible HTLV-1 Tax CEM-SS cells were resuspended in 10 ml of RPMI media and infected with either WT or IN- NL-NLuc virus in the presence or absence of 0.5 µg/ml dox. Before infection, viral supernatants were pretreated with 5 U/mL DNase I for 1 h at 37 °C to remove residual plasmid DNA. All IN- HIV-1 infections were supplemented with 20 µM Raltegravir to prevent revertant mutations. 10^6^ live cells were harvested at 1, 2, 3, 5, and 7 dpi and aliquoted to assay NLuc, and for DNA and RNA extraction.

For DNA analysis, cells were pelleted and washed three times in ice cold PBS and the cells incubated with DpnI (NEB) to remove any residual plasmid contamination. DNA was then extracted using DNA Miniprep Plus columns (Zymo Research) according to the manufacturer’s instructions.

For RNA analysis, cells were lysed in TRIzol (Thermo Fisher Scientific) and RNA harvested according to the manufacturer’s instructions, and DNAseI treated overnight. The RNA was then converted to cDNA using the High Capacity cDNA Reverse Transcription kit (Applied Biosystems).

All quantitative PCR reactions were performed in triplicate in a StepOnePlus or QuantStudio3 real-time PCR system according to the manufacturer’s instructions. Relative quantification of HIV-1 DNA levels was performed using the ΔΔCT method with β-Actin as an internal control (42). For experiments analyzing total HIV-1 DNA and RNA levels, HIV-1 DNA/cDNA was PCR amplified with a custom total HIV-1 TaqMan probe/primer set that amplifies the U5-gag region of HIV-1. β-Actin DNA was PCR amplified using a premade TaqMan probe/primer set, while β-Actin cDNA in RNA analyses was quantified using a separate Taqman probe/primer set amplifying across a splice junction of β-Actin. 2LTR DNA and RNA were similarly quantified using TaqMan primers/probes that amplify across the U5-U3 junction (43).

For Alu-LTR real time nested qPCR experiments, DNA was amplified using a modified version of the nested PCR approach described previously (28). Briefly, an initial non-saturating PCR using primers ALU1, ALU2, and L-HIV was performed using DNA isolated from HIV-1 infected cells. After the PCR products were purified using a PCR Kleen kit (Bio-Rad), a nested qPCR was performed using primers AA55M and L and the SYBR green master mix (Thermo Fisher Scientific).

### siRNA Knockdowns and sgRNA Knockouts

CD4+ 293T cells were transfected with 25 pmol of pooled siRNA (Origene, 3 siRNA per pool) with RNAiMax twice (on day 1 and 3), and infected on day 4 with WT or IN- NL-NLuc virus. Infected cells were assayed for NLuc activity 2 days post infection.

293T cells were transfected with pLentiCrispr v2 encoding the relevant sgRNA sequences using PEI. The cells were selected for puromycin resistance 2 days post-transfection, then single cell cloned.

The area around the NP220 sgRNA cut sites in the clones was sequenced to detect the expected deletion, and the expression of NP220 in the selected clones assayed by Western blot. Cells were then infected with VSV-G pseudotyped WT or IN- NL-NLucΔEnv reporter virus and assayed for NLuc activity 2 days post-infection.

Similarly, sgPML single cell clones were assayed for PML expression by Western blot. DNA from PML non-expressing cells was extracted, the PML regions of interest around the cut site amplified, cloned and sequenced.

### Western Blot Analyses

Cells were harvested and lysed in Laemmli buffer, sonicated and denatured at 95°C for 15 min. Lysates were subjected to electrophoresis on 4–20% SDS-polyacrylamide gels (Bio-Rad), transferred onto nitrocellulose membranes and then blocked in 5% milk in PBS + 0.1% Tween. Membranes were incubated in primary and secondary antibodies diluted in 5% milk in PBS + 0.1% Tween for 1 h each and then washed in PBS + 0.1% Tween. The membranes were incubated with a luminol-based enhanced chemiluminescent (ECL) substrate and signals were visualized using GeneSnap (Syngene).

### Southern Blot Analysis

Southern blots were performed as described previously (44). Briefly, DNA was extracted from CEM-SS or Tet-inducible HTLV-1 Tax CEM-SS cells that were infected with DNaseI-treated WT or IN- HIV-1 at 2dpi using a DNA Miniprep Plus kit (Zymo Research). Purified DNA was then digested with MscI, XhoI, and DpnI (NEB) overnight, recovered by standard ethanol DNA precipitation, then 10 µg of DNA was run on a 1% TBE gel. The gel was then soaked in denaturation solution (1.5 M NaCl, 0.5 M NaOH) to separate the double stranded DNA strands, before being washed in neutralization buffer (3 M NaCl, 0.5 M Tris-HCl pH 7.0). The DNA was transferred overnight onto a Gene Screen Plus nylon membrane (PerkinElmer) by capillary action, the membrane washed in 2xSSC, then UV-crosslinked in a Stratalinker 2400 at 3000 microjoules power (Stratagene). The membrane was then blocked in ExpressHyb hybridization solution (Clontech) for 1 hour.

HIV-1-specific DNA was amplified by PCR from the pNL4-3 plasmid using the primers: 5’-AGAAGAAATGATGACAGCATG-3’ and 5’-TGCCAGTTCTAGCTCTG-3’. The radiolabeled probe was random primed from this DNA amplicon using Prime-a-Gene Labelling System (Promega) and ^32^P-dCTP (Perkin Elmer). The probe was then denatured and hybridized onto the blocked membrane overnight at 60°C The membrane was then washed with solution 1 (2xSSC, 0.05% SDS) and solution 2 (1xSSC, 0.1% SDS) at 50°C for 1 hour each. Bands were visualized on film, and hybridization signals were quantified using a Typhoon phosphorimager (Amersham).

### ChIP-qPCR

10^7^ Tet-inducible HTLV-1 Tax CEM-SS cells were resuspended in 10 ml of RPMI media and infected with either WT or IN- HIV-1 virus. 48 h post-infection, cells were rinsed twice with PBS and cross-linked with 1% formaldehyde for 20 min at 25°C, quenched in 0.125 M glycine for 5 min, and lysed in ChIP lysis buffer (50 mM Tris·HCl, pH 8.0, 1% sodium dodecyl sulfate, 10 mM EDTA). Cell lysates were then sonicated with a Fisher Sonic Dismembrator 60 (output 4.5, 20s pulse repeated 6 times on ice with 40 sec between each sonication). The supernatant containing sonicated chromatin was pre-cleared by the addition of magnetic Protein G Dynabeads (ThermoFisher) that had been pretreated with denatured salmon sperm DNA (Invitrogen). The magnetic beads were removed, and the sonicated chromatin was immunoprecipitated overnight at 4°C using 2.5 µg antibody in ChIP dilution buffer (16.7 mM Tris-HCL pH 8.0, 1% Triton X-100, 0.01% SDS, 150 mM NaCl, 1.2 mM EDTA). 5% of the sonicated chromatin was stored as input DNA without further treatment until the reverse crosslinking step. The next day, the incubated chromatin-antibody mixture was incubated with the pretreated Dynabeads for 2 hours at 4°C, and then washed with ChIP low-salt buffer (20 mM Tris-HCL pH 8.0, 1% Triton X-100, 0.1% SDS, 150 mM NaCl, 2 mM EDTA), ChIP high-salt buffer (20 mM Tris-HCL pH 8.0, 1% Triton X-100, 0.1% SDS, 500 mM NaCl, 2 mM EDTA), ChIP LiCl buffer (10 mM Tris-HCL pH 8.0, 1% NP-40, 250 mM LiCl, 1 mM EDTA, 1% Na Deoxycholate) and TE buffer (10 mM Tris HCL pH 8.0, 1 mM EDTA). Protein-DNA complexes were eluted from the beads with an Elution buffer (0.1 M NaHCO_3_, 1% SDS), then de-crosslinked by incubating at 65°C for 16 hours and 95°C for 15 mins, then proteins removed by adding proteinase K at 50°C for 3 hours. DNA was then purified using a DNA Miniprep Plus kit (Zymo), then digested with DpnI (NEB) to remove any plasmid contamination, before being used for qPCR analysis using primers that amplify U5-R on HIV-1 and the SYBR green master mix (Thermo Fisher Scientific). ΔΔCt was calculated relative to total histone H3 levels and expressed as fold change relative to cells not expressing a viral protein and infected with WT HIV-1 in the bar graphs.

### Quantification and Statistical Analysis

Descriptive statistics were performed with GraphPad Prism 8 (GraphPad Software, US). The size of each study or number of replicates, along with the statistical tests performed, can be found in figure legends. Numerical data are presented as the mean ± standard deviation.

## Figure Legends

**Figure S1:**
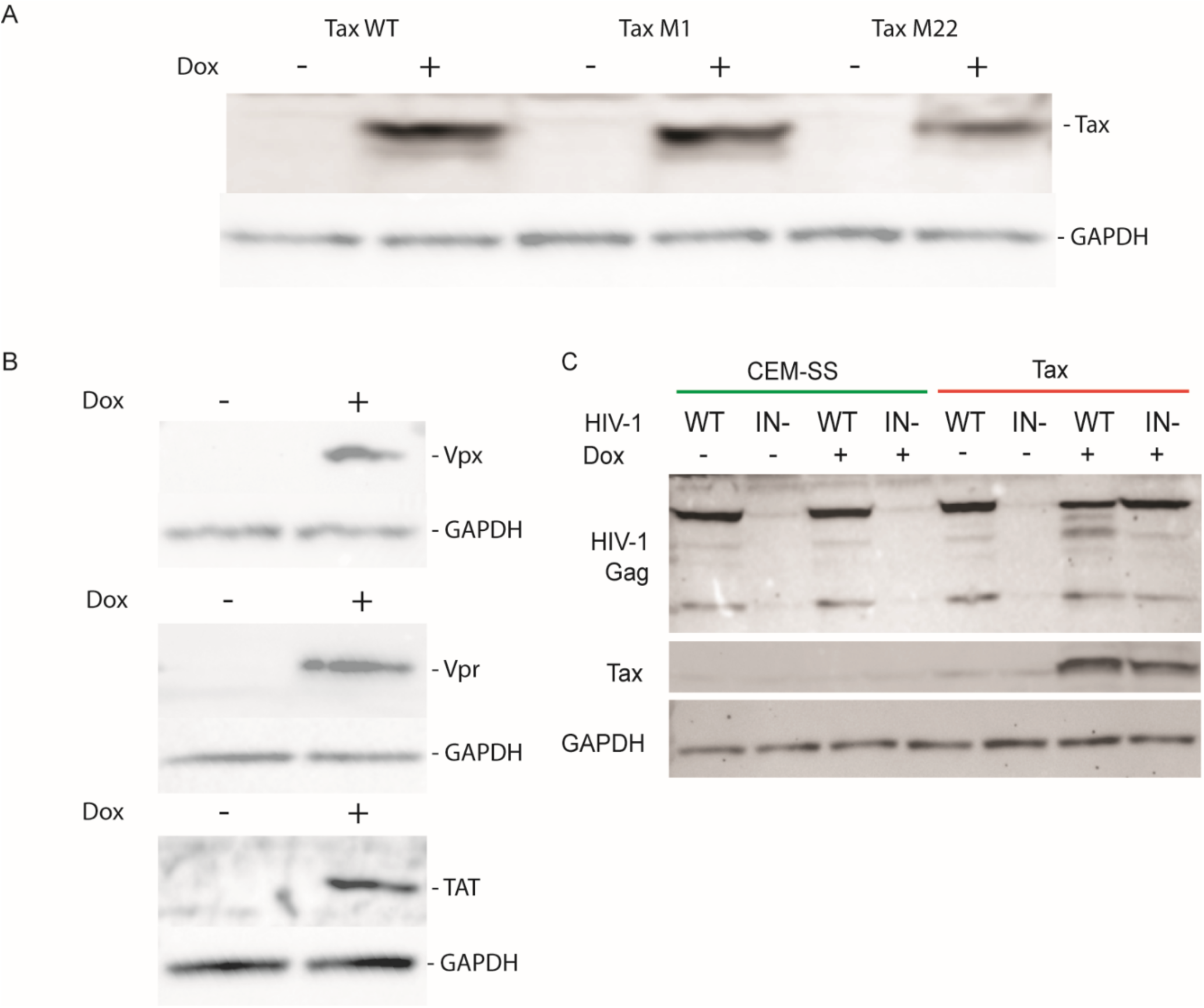
Dox-inducible expression of Tax but not Vpx, Vpr or Tat rescues the expression of HIV-1 gag from unintegrated HIV-1 episomes. **A)** Single cell clones of CEM-SS cells transduced with a Tet-inducible lentivector expressing HTLV-1 Tax or the indicated Tax mutants in the presence or absence of 0.5 µg/ml Dox. **B)** Similar to **A** but cells were transduced with Tet-inducible lentivectors expressing HIV-2 Vpx, HIV-1 Vpr, or HIV-1 Tat. **C)** WT CEM-SS cells and tet-inducible, Tax expressing CEM-SS cells were infected with WT or IN- HIV-1 in the presence or absence of Dox and probed on a Western blot for the indicated proteins.

**Figure S2:**
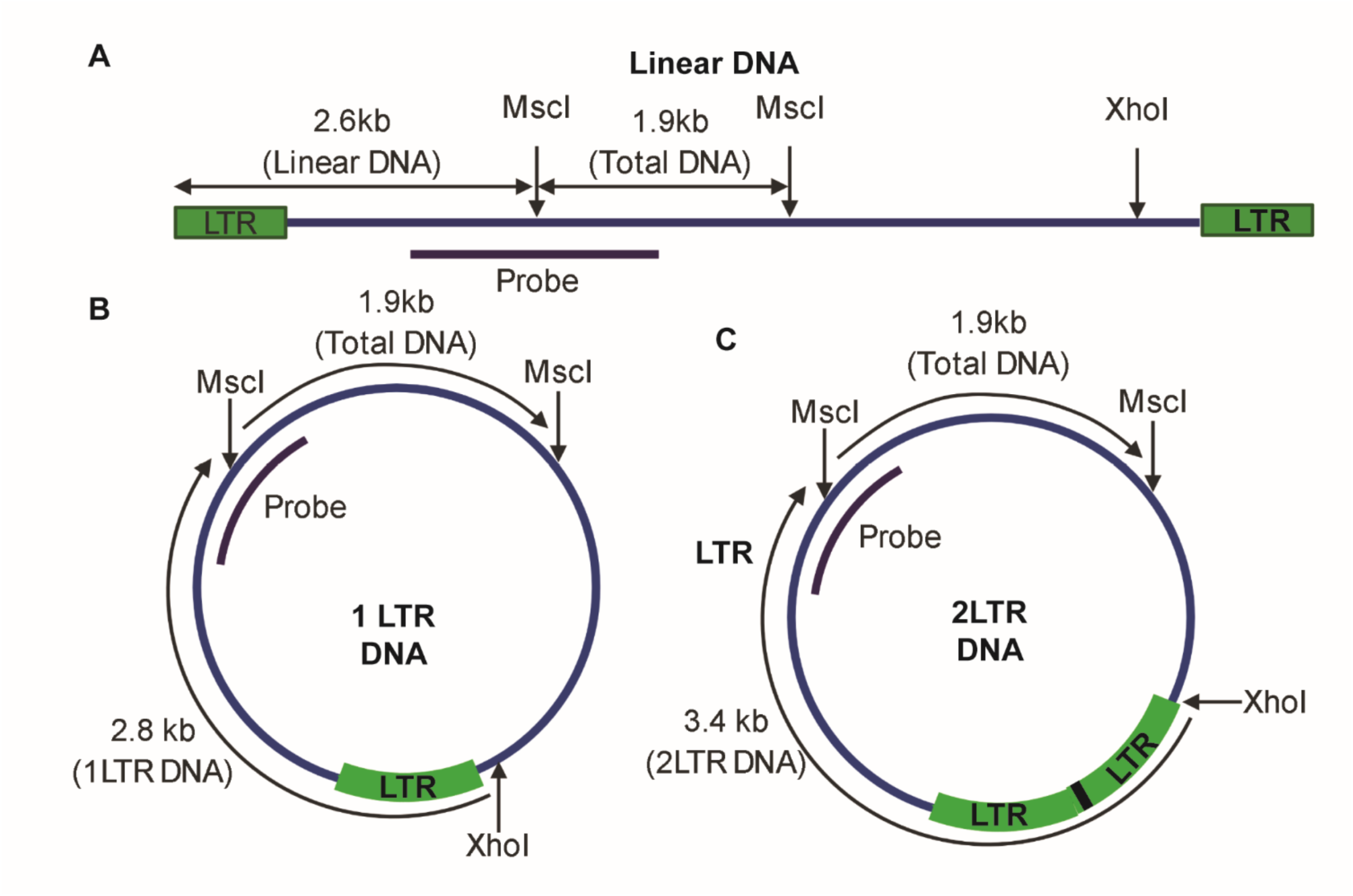
Southern blot restriction enzyme and probe binding schematic. DNA from HIV-1 infected cells was digested with MscI and XhoI (and DpnI to remove residual plasmid contamination) overnight. This cuts HIV-1 DNA at the indicated points, and these DNA fragments were then separated by gel electrophoresis. A P^32^ radio-labelled DNA probe that spans an MscI cut site was used to detect the DNA fragments (**A-C**, approximate probe binding in purple). This probe detects a 1.9kb DNA fragment that is released by all HIV-1 DNA forms (**A-C**), and also a 2.6 kb fragment released by unintegrated linear HIV-1 DNA (**A**), a 2.8 kb fragment released by 1LTR circle DNA (**B**), and a 3.4 kb fragment released by 2LTR circle DNA (**C**). Integrated HIV-1 DNA only produces the 1.9 kb fragment.

**Figure S3:**
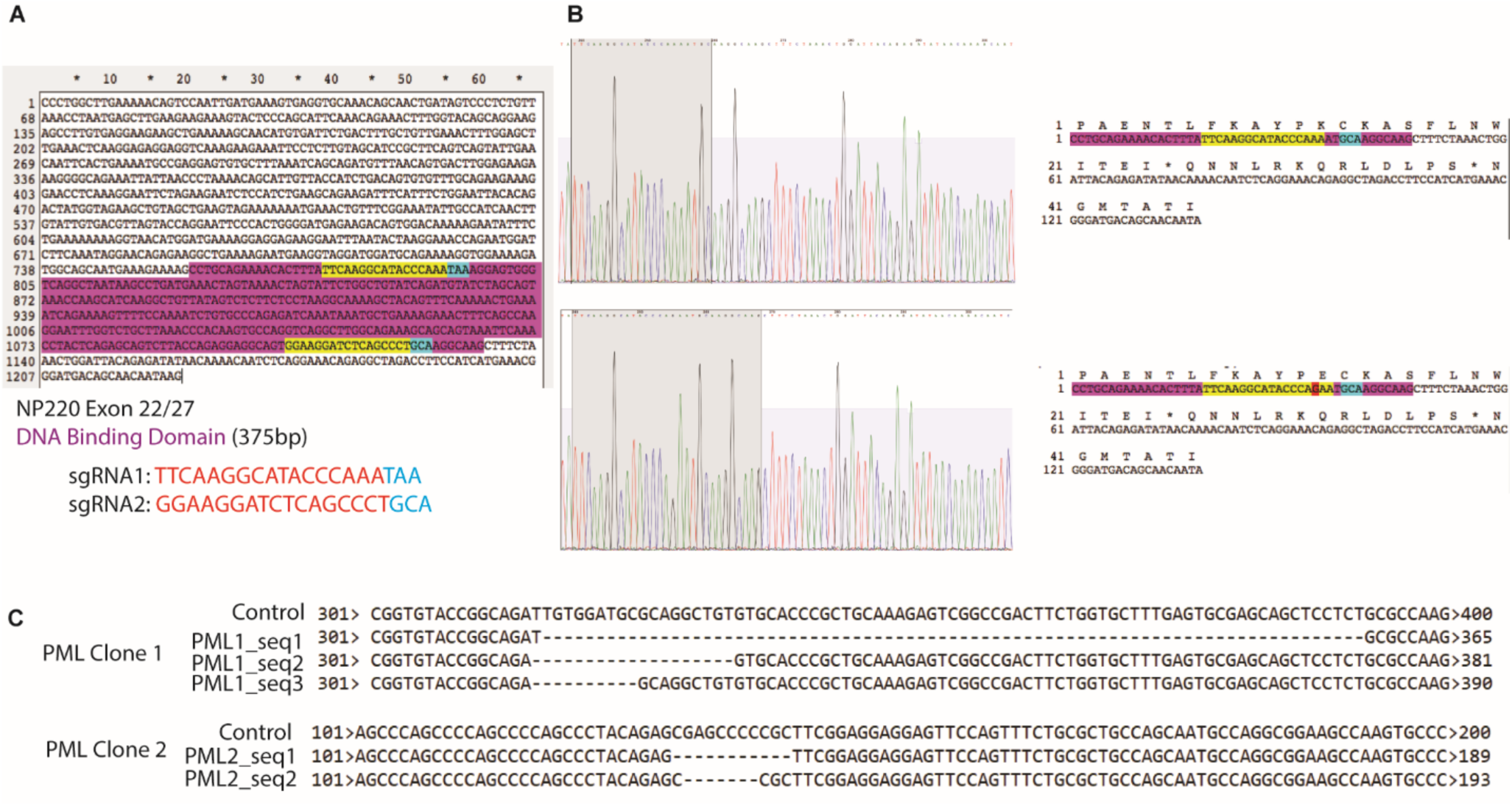
NP220 and PML knock-out 293T cell clones were generated using Crispr/Cas. **A)** The NP220 DNA binding domain on exon 22 (highlighted in purple) was targeted for excision using two sgRNAs, highlighted in yellow and blue. The yellow/blue boundary is the predicted Cas9 cleavage site. **B)** DNA sequences of clones of 293T cells transfected with the two sgRNAs showing excision of the NP220 DNA binding domain and the introduction of premature stop codons. **C)** PML knockouts were generated by transfection of 293T cells with lentiCrisprv2 encoding either sgPML1 or sgPML2. Single cell clones were isolated after puromycin selection, the targeted region of the PML gene was PCR amplified, cloned and sequenced. PML clone 1 (sgPML1) has 10, 19 and 77 bp deletions, while PML clone 2 (sgPML2) has 7 and 11 bp deletions, all of which are predicted to introduce frame shift mutations.

**Table S1.**
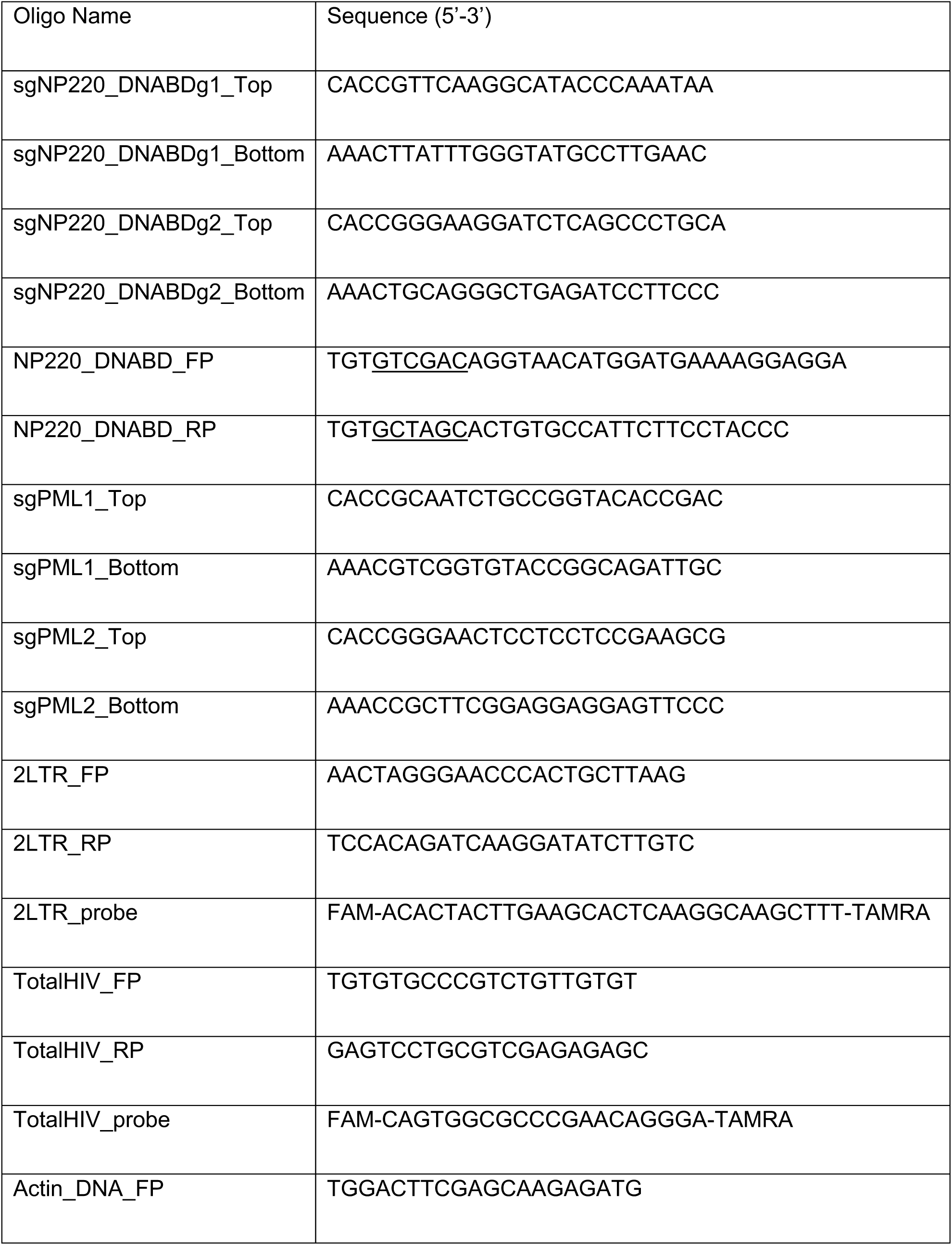
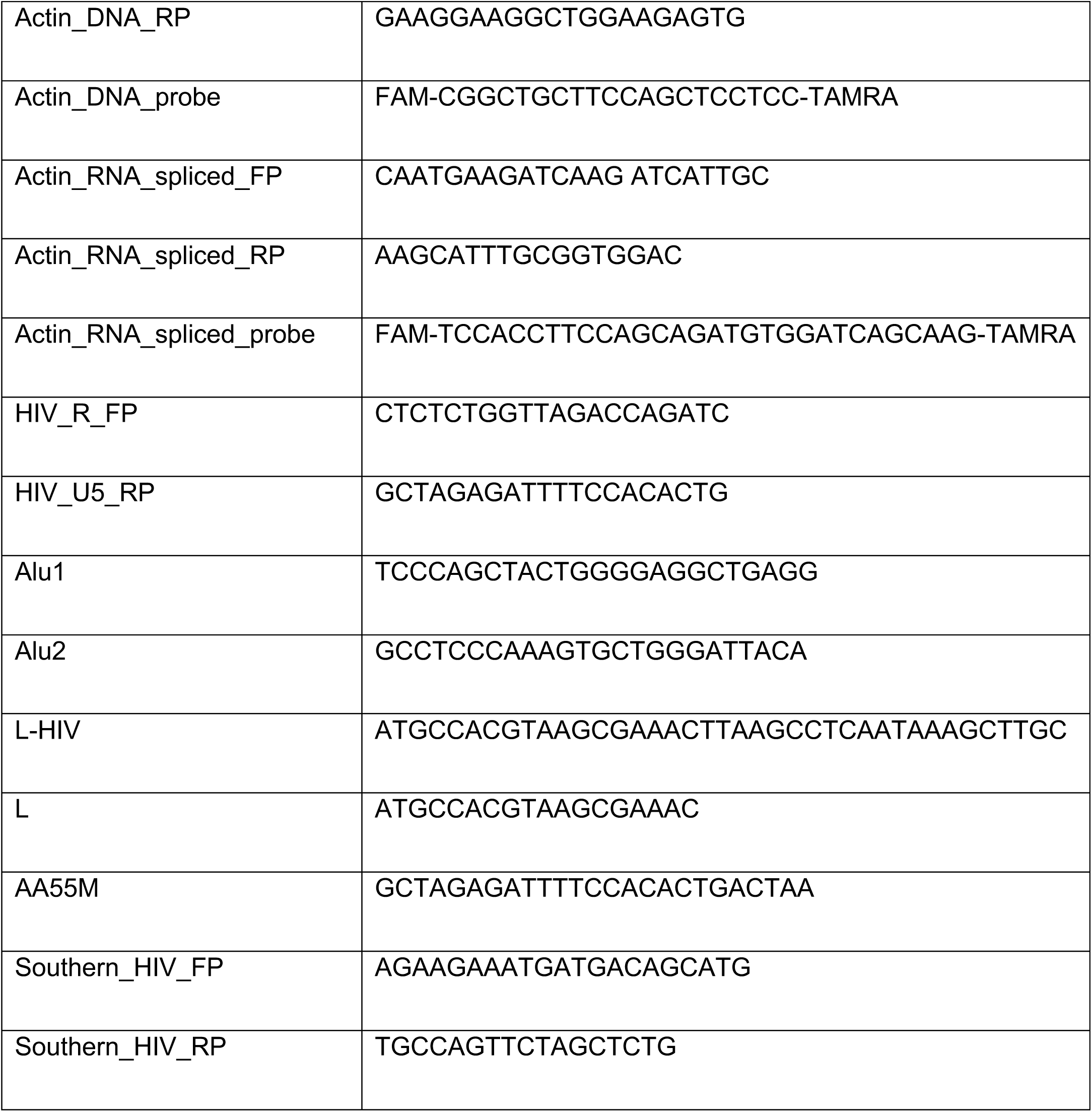
Oligonucleotide sequences. Related to the STAR Methods.

